# Spatial Mapping of Mobile Genetic Elements and their Cognate Hosts in Complex Microbiomes

**DOI:** 10.1101/2023.06.09.544291

**Authors:** Benjamin Grodner, Hao Shi, Owen Farchione, Albert C. Vill, Ioannis Ntekas, Peter J. Diebold, Warren R. Zipfel, Ilana L. Brito, Iwijn De Vlaminck

## Abstract

The frequent exchange of mobile genetic elements (MGEs) between bacteria accelerates the spread of functional traits, including antimicrobial resistance, within the human microbiome. Yet, progress in understanding these intricate processes has been hindered by the lack of tools to map the spatial spread of MGEs in complex microbial communities, and to associate MGEs to their bacterial hosts. To overcome this challenge, we present an imaging approach that pairs single molecule DNA Fluorescence In Situ Hybridization (FISH) with multiplexed ribosomal RNA FISH, thereby enabling the simultaneous visualization of both MGEs and host bacterial taxa. We used this methodology to spatially map bacteriophage and antimicrobial resistance (AMR) plasmids in human oral biofilms, and we studied the heterogeneity in their spatial distributions and demonstrated the ability to identify their host taxa. Our data revealed distinct clusters of both AMR plasmids and prophage, coinciding with densely packed regions of host bacteria in the biofilm. These results suggest the existence of specialized niches that maintain MGEs within the community, possibly acting as local hotspots for horizontal gene transfer. The methods introduced here can help advance the study of MGE ecology and address pressing questions regarding antimicrobial resistance and phage therapy.

## INTRODUCTION

Understanding the complex biology of mobile genetic elements (MGEs) is crucial for manipulating microbiomes and improving the treatment of microbiome-associated diseases. MGEs carried on plasmids can confer adaptive traits, including antimicrobial resistance (AMR) and virulence, to host bacteria, while bacteriophages can drastically alter the structure of microbiomes.^1–3^ The host range of MGEs varies widely — some have a broad host range, while others are restricted to a single strain or species. This host range is consequential; for example, the host range of bacteriophages can impact their utility for precision microbiome manipulation or infection treatment.^4^ Similarly, the host range of AMR plasmids may inform the extent to which a microbiome can act as a reservoir for AMR traits.^5,6^

Despite the centrality of MGEs to microbial ecology, basic facts about the mechanisms of the spatial spread of MGEs within natural communities remain unknown. This knowledge gap largely stems from a lack of tools to examine the mobile gene pool in situ and to directly establish MGE-host associations.^7^ Metagenomic sequencing is the most common tool used to study the repertoire of MGEs in microbiomes, but metagenomic sequencing struggles to associate MGEs with host bacteria, and does not retain spatial information.

In this study, we introduce an imaging-based approach that integrates single-molecule DNA Fluorescence In Situ Hybridization (FISH) and highly multiplexed rRNA-FISH to map MGEs and their cognate bacterial hosts at the resolution of a single bacterial cell. We show that this method enables to study the heterogeneity in the spatial distribution of MGEs within biofilms, and to establish links between MGEs and their hosts in complex structured microbiomes. We developed this method for confocal microscopy with spectral detection to situate MGEs in three dimensions within dense biofilms and to enable simultaneous highly multiplexed identification of bacterial taxa. We first assessed and optimized single molecule DNA FISH techniques based on *in situ* signal amplification to ensure sensitive and specific detection of target DNA within individual bacterial cells via confocal microscopy. Next, we developed a semi-automated image analysis pipeline to detect MGE spots and segment bacterial cells. We then applied this methodology to examine the spatial spread of AMR-carrying plasmids and prophage in human oral plaque biofilms. We demonstrated the ability to establish MGE-host associations, and we found that both bacterial taxa and their MGEs exhibit intricate spatial structure, forming clusters within plaque biofilms on the order of 10-100μm. This spatial heterogeneity implies the existence of diverse microscale niches of MGEs in dense biofilms and, potentially, taxonomic and physical barriers for horizontal gene transfer.

## RESULTS

### Optimization of single molecule MGE FISH

We used *Escherichia coli* transformed with pJKR-H-tetR plasmids encoding an inducible *GFP* gene as a model system to assess and optimize MGE-FISH on a confocal microscope (**Fig. 1a**).^8^ We designed FISH probes for the non-coding strand of the *GFP* gene, used non-transformed *E. coli* as a negative control, and tested six different FISH protocols. Initial attempts using single and ten encoding probes yielded little to no separation between the signal in the plasmid and control samples (**Fig. 1b**, *rows 1&2*). This was expected given the photon noise and losses inherent to confocal microscopy as compared to a wide field microscope.^9,10^ We next implemented two enzyme-free amplification methods to increase the signal.^11,12^ Branched amplification yielded a higher true positive signal, albeit accompanied with a high background signal in the negative control (**Fig. 1b**, *row 3*). Hybridization Chain Reaction (HCR) similarly enhanced the signal at the expense of a high background in the control (**Fig. 1b**, *row 4*). In order to improve specificity, we adopted a “split” HCR method and used heat-denatured DNA and non-fluorescent “helper probes’’ to stabilize the DNA.^13,14^ This resulted in a significant reduction of the signal in the negative control (**Fig. 1b**, *row 5)*. Last, to address autofluorescence in oral biofilms (as detailed below), we applied a gel embedding and clearing technique, in which nucleic acids in the sample are covalently anchored to a polyacrylamide gel, followed by clearing of proteins and lipids.^15,16^ That method led to a high specificity of MGE detection (false positive rate < 0.01) but a relatively low sensitivity (true positive rate = 0.39). We suggest that this limited sensitivity is a result of tight packing of the transcriptionally repressed GFP gene, limiting accessibility, as detailed previously and as confirmed by our experiments with a phage infection model described below.^17–19^ We applied the final optimized method in conjunction with super-resolution Airyscan imaging to examine the subcellular localization of plasmid-encoded *GFP* in *E. coli* cells. We found that the plasmid density is approximately 50% higher on average at the poles compared to the center (**Fig. S1a**,**b**), in line with previous reports that plasmids have limited capacity to diffuse through the nucleoid at the cell center and tend to cluster at cell poles.^20,21^

**Figure 1.**
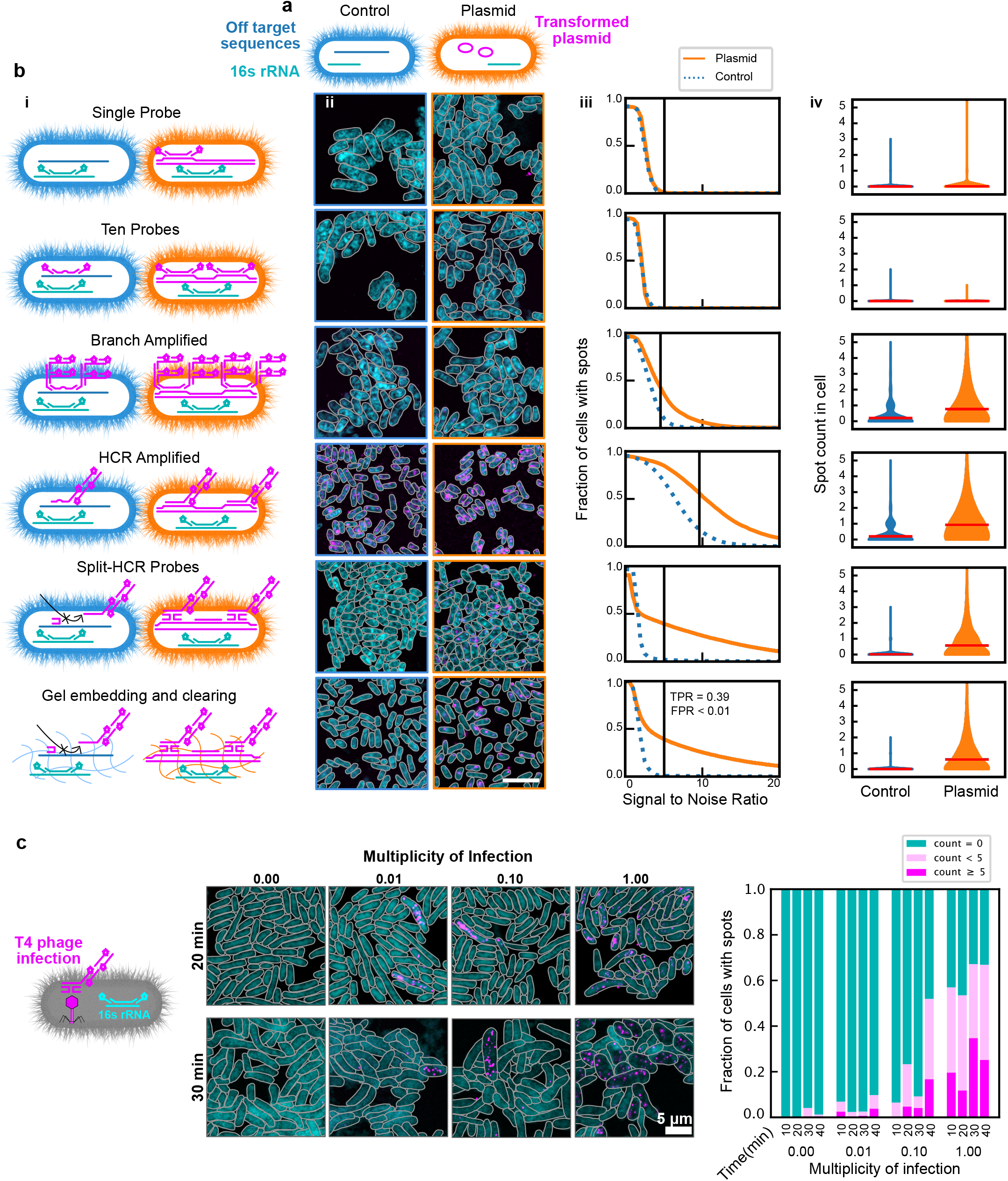
Single-molecule MGE FISH. **a** Diagram of *E. coli* model GFP plasmid system used to optimize smFISH. **b** *Panel i*: diagrams of different methods implemented. Blue cells on the left are wild type and orange cells on the right are transformed with the plasmid. After the first row, two encoding probes are shown to represent ten encoding probes in all cases. *Panel ii*: representative images for each method alteration. Scale bar is 5μm. *Panel iii*: fraction of cells with spots for control and plasmid images as a function of signal to noise ratio. Black vertical line indicates the selected SNR threshold. TPR: true positive rate; FPR: false positive rate (at the threshold). *Panel iv*: histograms for the number of spots in each cell. Width indicates the frequency of the spot count value. Horizontal red bars indicate mean spot count. **c** *Le*.: Diagram of MGE-FISH staining of *E. coli* infected by T4 Phage. *Center*: example images for four multiplicities of infection 20 minutes and 30 minutes after introducing phage to the culture. *Right*: results of manual counting to classify cells into groups based on the number of MGE-FISH spots.

### Visualizing phage infection

Building on the optimized MGE-FISH method (**Fig. 1b**, *row 6*), we turned our attention to visualizing T4 phage infection of *E. coli*. We staged infections at four multiplicities of infection (MOI 0, 0.01, 0.1, and 1), and took snapshots every ten minutes over a 40-minute period (**Fig. 1c, Fig. S1c**). We designed FISH probes targeting the non-coding strand of the *gp34* gene, which encodes a tail fiber protein and quantified cells with 5 or more MGE spots, less than 5 spots, and no spots (**Fig. 1c, Fig. S1d**). For non-infected controls (MOI 0), the fraction of cells with phage detected was 0.015 (8800 cells, 3 fields of view), which gives the false positive rate. No cells in the MOI 0 control had more than 5 spots, which gave us confidence that the striking signal from cells with high spot count in MOI 0.01, 0.1, and 1 was specific to phage infection. We predicted the fraction of infected cells to be 0.00, 0.01, 0.10, and 0.73, for MOI 0, 0.01, 0.1 and 1 respectively (Poisson probability mass function). This was close to the observed fraction of cells with phage spots at 20 minutes: 0.00, 0.02, 0.23, and 0.53. This indicates much higher sensitivity than what we observed in the *GFP* plasmid experiment (**Fig. 1b**, *row 6*). We suggest that the actively replicating *gp34* gene is more accessible to FISH probes than the transformed, unexpressed *GFP* gene in the plasmid experiment.

T4 phage infecting *E. coli* in LB media has a reported average latent period lasting 18 minutes, end of lysis at 36 minutes, and a burst count of 110.^22^ We observed MGE-FISH spots within 10 minutes of phage introduction, which indicates that we are visualizing replicated phage genetic material before disruption of the cell membrane. At 20 minutes, cells with high phage count were often physically longer in length than uninfected cells, suggesting bacterial growth with stalled division near the end of the latent period. Our results match previous findings that burst sizes for T4 phage increase with increased bacterial growth rate due to large cell volumes delaying full lysis.^22,23^ We observed a dramatic increase in the fraction of infected cells for MOI 0.01 and 0.1 at 40 minutes. This corresponds to the expected lysis time and the adsorption of new phage to uninfected cells. At 30 and 40 minutes, many cells with a high phage count had a low 16S rRNA signal and increased width and length compared to uninfected cells (**Fig. 1c, Fig. S1c**). We suggest that these cells with high phage count and low 16S rRNA intensity have been fully lysed, meaning that MGE-FISH can be used to stain encapsulated phage particles, as has been suggested previously.^24^ We also observed a small fraction of infected cells with a low 16S rRNA signal in the center of the cell and a high signal at the poles (**Fig. S1d**, *middle*), which we suggest are infected cells that experience cytoplasmic condensation due to membrane damage.^25^ Overall, these data and observations match the expected progression of a T4 phage infection course, and show the value of MGE-FISH imaging to generate novel insights even in a well studied system.

### Mapping MGEs in oral plaque biofilms at high specificity

Next, we evaluated the ability of our MGE-FISH method to visualize the spatial distribution of MGEs in human oral plaque biofilms. To this end, we collected oral plaque biofilms from two healthy volunteers (A and B) and performed shotgun metagenomic sequencing on a portion of each sample, reserving the rest for imaging (**Fig. 2a**). As an initial controlled test of the method (**Fig. 1b**, *row 6*), we stained for the *GFP* gene in samples that contained mixtures of plaque and *GFP*-transformed *E. coli* (**Fig. 2b**) and demonstrated that the specificity remained high in plaque.

**Figure 2.**
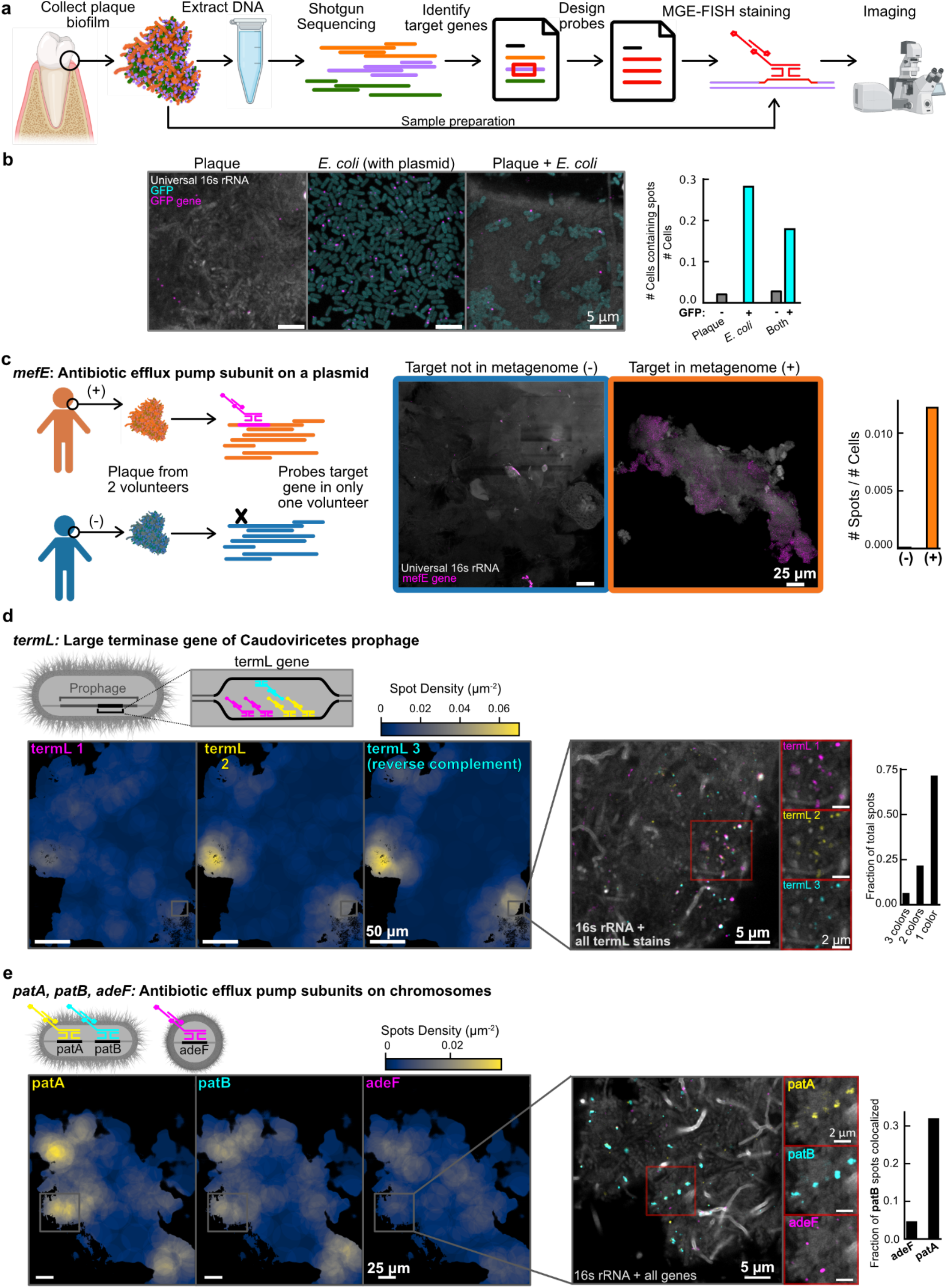
MGE-FISH in human oral plaque. **a** Diagram of the workflow to apply MGE-FISH in oral plaque biofilms. **b** *Left*: Example images of standard plaque, transformed *E. coli* expressing GFP, and the combination of both plaque and *E. coli*. All samples were stained for the *GFP* gene using MGE-FISH. *Right*: association of MGE-FISH signal with GFP cells and non-GFP cells in each sample. **c** *Left*: Diagram of two-volunteer control experiment. *Center*: example images of plaque samples from each volunteer stained for the *mefE* gene. *Right*: measurement of relative spot count for each volunteer. **d** *Top left*: diagram showing the multicolor approach used to stain the gene *termL*. Bottom leM: example FOV plotted as density maps for each color of *termL* probes. *Inset 1*: zoomed region of the plaque overlaid with all colors of *termL* stain. *Inset 2*: zoomed region of plaque split into each color of *termL* probes. *Right*: measurements of *termL* color colocalization normalized as the fraction of total spots. **e** *Top left*: diagram showing the multicolor approach used to simultaneously stain the genes *patA, patB*, and *adeF. Bottom left*: example FOV plotted as density maps for each gene. *Inset 1*: zoomed region of the plaque overlaid with all colors. *Inset 2*: zoomed region of plaque split by gene. *Right*: measurement of colocalization of *patB* spots with each other gene normalized as the fraction of *patB* spots colocalized.

Via metagenomic analysis, we identified *mefE*, an AMR gene located on a plasmid and encoding an antibiotic efflux pump, in the plaque of volunteer A but not volunteer B (**Fig. 2c**). Our MGE-FISH method confirmed the prediction from metagenomic analysis; we measured 0.012 and 0.000 *mefE* spots per cell in volunteers A and B respectively (**Fig. 2c**). Furthermore, we demonstrated that there was positive spatial autocorrelation of *mefE* spots in volunteer A (Moran’s I = 0.015, p=0.005, **Fig. S2a**), suggesting that the process underlying the distribution of plasmids was non-random, while the spots in volunteer B were randomly distributed (Moran’s I = 0, p=0.259). These results showed that MGE-FISH is effective to visualize MGEs in plaque. The spatial clustering of this AMR plasmid suggests that there are regions within the biofilm that promote localized spread of the AMR plasmid, perhaps through vertical transfer during replication of host cells, or through horizontal transfer between neighbors in the region.^26^

In the plaque, we observed off-target signals as bright patches and dispersed large spots, likely due to nonspecific binding of probes to food particles or debris. To mitigate this issue, we implemented gel embedding and clearing for reduced off-target binding.^24,32^ To test the efficacy of gel embedding and clearing, we used orthogonal FISH probes, designed to not target any sequence in the plaque. We observed a dramatic reduction in off-target signal after gel embedding and clearing (**Fig. S2b**,**c**), and therefore used this in all subsequent experiments on plaque.

We next mapped a natural lysogenic bacteriophage (prophage) in plaque to study its spatial distribution. In volunteer B, we identified a T7-like prophage via metagenomic analysis and developed probes targeting its *capsB* gene, which encodes the minor capsid protein. In these experiments, we used two negative controls to assess off-target binding: one with no probe and one with orthogonal probes. Both controls displayed minimal off-target signal (**Fig. S3a**), and we could set an area threshold on spots to further filter out off-target signals based on the spot size. *CapsB* spots clustered spatially, coinciding with long, rod-shaped bacteria. The spatial clustering of this phage is likely due to a limited host range; in the metagenomic analysis this prophage was binned with *Corynebacterium*, a long rod shaped bacteria that forms spatial clusters.^27^ Large clusters (∼100 μm) of host bacteria may result in localized hotspots of prophage spread in a biofilm (**Fig. S3b**).

In order to further test the robustness of MGE-FISH in plaque, we then proceeded to label another phage gene in three different colors simultaneously. We identified a highly prevalent prophage of the class *Caudoviricetes* with a large terminase gene, *termL*, and were able to design a large set of FISH probes. We divided the probes into three groups, each labeled with a different color. We mapped the large-scale distribution (∼25 μm) of spots in each color and found that they formed similar patterns, as expected (**Fig. 2d**). We also demonstrated that different color spots colocalized with each other at the micron scale. Similar to the previous prophage, this prophage also formed isolated spatial clusters, suggesting spatial restriction of host bacteria within plaque biofilms. While dense clusters of host cells could result in rapid transfer of a lytic phage within the cluster, the spatial isolation of different host clusters may limit the global spread of infection, with the intervening non-host cells acting as a barrier to phage transfer.

In addition to MGEs, we also tested the possibility to visualize genes located on bacterial genomes. Using metagenomic analysis, we identified three non-plasmid AMR genes. Genes *patA* and *patB*, subunits of an antibiotic efflux pump, were from the same Metagenomic Assembled Genome (MAG) and had nearly identical coverage values, so we expected them to spatially colocalize. We found another antibiotic efflux pump subunit, *adeF*, in a different MAG (**Fig. 2e**). At the large scale (∼25μm), *patA* and *patB* had similar density patterns, while *adeF* had a distinct pattern, as expected. At the micron scale, MGE-FISH staining for these three genes showed that 32% of *patB* spots colocalized with *patA*, while only 5% of *patB* spots colocalized with *adeF*. The difference in large scale spatial distribution between *patA/B* and *adeF* indicates that cells carrying these AMR genes occupy different spatial niches. Identifying spatial niches for AMR genes within biofilms via MGE-FISH can help gain understanding of the maintenance and spread of AMR.

### Combined taxonomic mapping and MGE mapping

We next strived to overlay MGE biofilm maps with taxonomic identity maps to associate MGEs with their host taxa. To start, we measured the taxonomic association of a highly prevalent prophage of class *Caudoviricetes*, for which the metagenomic data hinted at a strong taxonomic association with *Veillonella* (**Fig. 3a**).^28–31^ We used rRNA FISH to stain five common oral genera, *Veillonella, Streptococcus, Corynebacterium, Lautropia*, and *Neisseria*, each with a different fluorophore, and we used MGE-FISH to stain the *termL* gene of the active prophage with a sixth fluorophore (**Fig. 3b**). The *termL* gene and *Veillonella* showed striking colocalization, mirroring the prediction from metagenomic assembly (**Fig. 3c**). We quantified the fraction of *termL* spots within 0.5 μm of each species and compared the observed values to simulations of randomly distributed spots. *Veillonella* displayed by a large margin the highest spatial association considerably above random (Z score = 7.7, p ≤ 0.01, **Fig. 3d**). The fraction of *termL* spots associated with *Veillonella* was 0.39, while the fraction *termL* associated with each other genus was very low (∼0.01). These results clearly demonstrated our ability to determine MGE host taxonomy in plaque biofilms by concurrently mapping taxa identity and MGEs.

**Figure 3.**
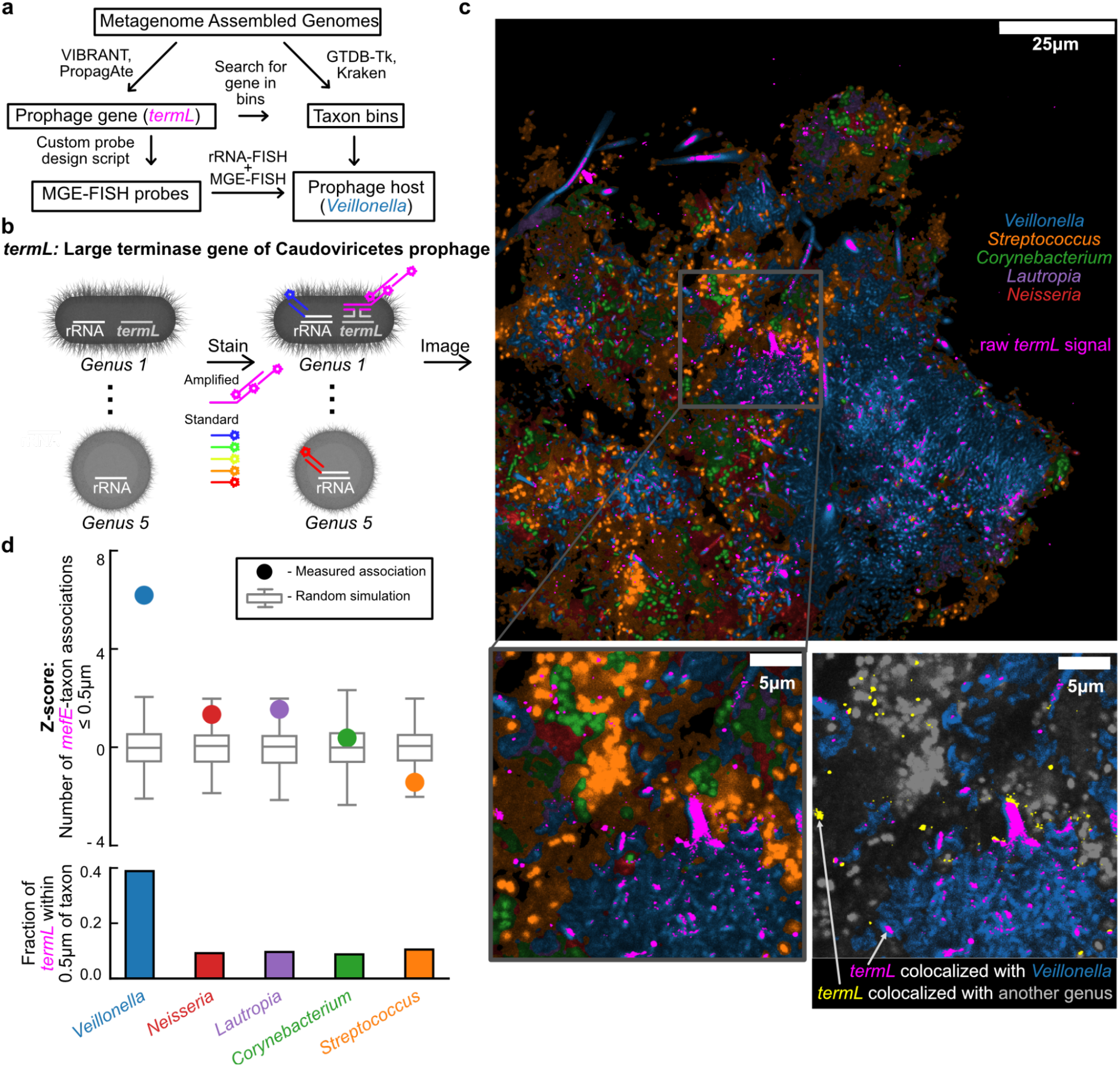
Combined MGE and taxonomic mapping. **a** Workflow for orthogonal prophage host association predictions via metagenomic sequencing analysis or MGE-FISH with rRNA-FISH taxon mapping. **b** Diagram showing simultaneous taxon mapping and MGE mapping. **c** *Top*: Bacterial genera classified by rRNA-FISH overlaid with the raw signal from MGE-FISH on *termL. Bottom left*: zoomed region of rRNA-FISH overlaid with MGE-FISH. *Bottom right*: zoomed region showing only *Veillonella* (blue) and *termL* (magenta and yellow) in color, while all other cells are grayscale. The arrows indicate examples of *termL* signal colocalized with *Veillonella* in magenta, and *termL* signal colocalized with another genus in yellow. **d** *Top*: z-scores for the number of associations between *termL* and each genus (circles) compared to simulation of random distributions of the same spots (boxplots, 1000 simulations). *Bottom*: fraction of *termL* spots associated with each taxon. Association of a cell with a spot is defined as separation less than or equal to 0.5μm.

Subsequently, we sought to identify the host of an AMR plasmid for which metagenomic binning did not yield a candidate host. The CARD database identified *mefE* on the plasmid as a subunit of a major-facilitator-superfamily antibiotic efflux pump.^32,33^ Because metagenomic sequencing data couldn’t predict host association, we broadened our target panel for taxonomic mapping by employing HIPR-FISH, a method which uses combinatorial spectral barcoding to map taxa. We developed a target panel of 18 genera that are highly abundant and prevalent in human plaque. We designed a HiPR-FISH probe panel using a 5-fluorophore combinatorial barcoding scheme, whereby each fluorophore represents a binary bit, providing 31 possible barcodes (2^5^-1 = 31 possible barcodes).^27,34^ The fluorophore for the MGE was spectrally distinct from those of HiPR-FISH, enabling simultaneous implementation of both methods (**Fig. 4a**).

**Figure 4.**
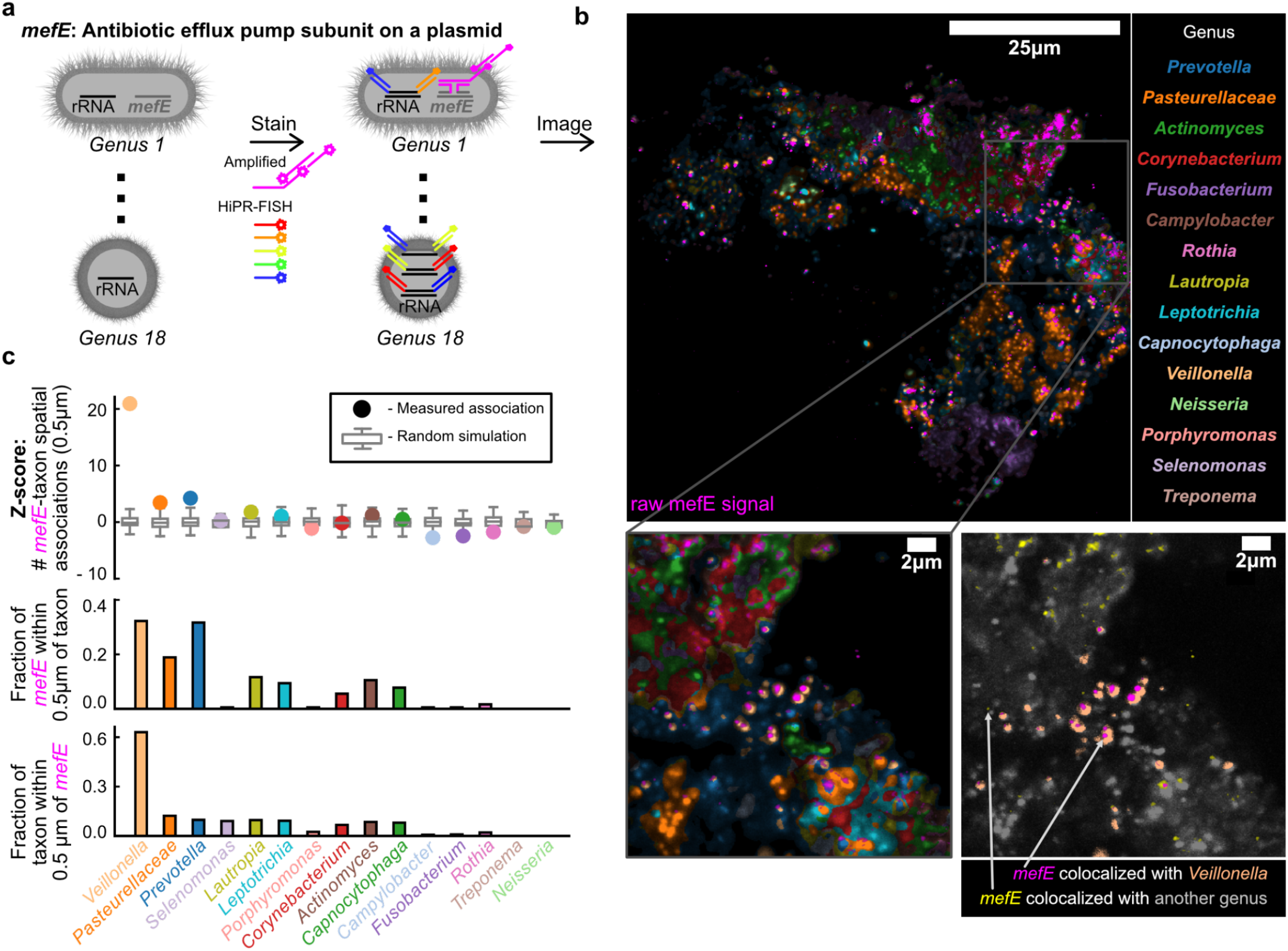
Identification of the host taxon of an AMR plasmid. **a** Diagram illustrating simultaneous HiPR-FISH combinatorial spectral barcoding and MGE-FISH. **b** *Top left*: Overlay of bacterial genera classified by HiPR-FISH and *mefE* mapped by MGE-FISH. *Top right*: taxon color legend for HiPR-FISH classification. *Bottom left*: zoomed region of HiPR-FISH overlaid with MGE-FISH. *Bottom right*: zoomed region showing only *Veillonella* (peach) and *mefE* (magenta and yellow) in color, while all other cells are grayscale. The arrows indicate examples of *mefE* signal colocalized with *Veillonella* in magenta, and *mefE* signal colocalized with another genus in yellow. **c** *Top:* z-scores for the number of associations between *mefE* and each genus (circles) compared to simulation of random distributions of the same spots (boxplots, 1000 simulations). *Middle*: fraction of *mefE* spots associated with each genus. *Bottom*: fraction of each genus associated with *mefE* spots. Association of a cell with a spot is defined as separation less than or equal to 0.5μm.

Using integrated HiPR-FISH and MGE mapping, we observed that *mefE*, like *termL*, strongly associated with *Veillonella* at a range of 0.5μm (Z-score = 20.9, p ≤ 0.01, **Fig. 4b**,**c**). This association was quantified from two perspectives: from *mefE*’s point of view, 32% of *mefE* spots were within 0.5μm of a *Veillonella* cell; from *Veillonella’s* perspective, 63% of its cells were within 1μm of a *mefE* spot (p ≤ 0.001). The visually and quantitatively prominent association of *mefE* with *Veillonella* suggests that *Veillonella* is the host for the *mefE* plasmid. This was further corroborated by the fact that mefE tended to localize within cauliflower structures, which are known to be formed with *Veillonella*.^27^ The majority of *Veillonella* cells carry this plasmid, and it is possible that the dense packing in cauliflower structures facilitates promiscuous HGT among *Veillonella* cells. Alternatively, the *Veillonella* cells observed might be descendants of a single strain carrying the plasmid, the plasmid being preserved over time due to this strain’s dominance in the niche. We also found that *mefE* is associated with *Pasteurellaceae* and *Prevotella* more frequently than simulations predict. However, we observed that *Pasteurellaceae* and *Prevotella* physically associate with *Veillonella*. The plasmid’s transfer from *Veillonella* to Pasteurellaceae or *Prevotella* seems unlikely given the significant phylogenetic distance between them, as HGT usually occurs between closely related species. All in all, these experiments constitute a demonstration of the use of DNA FISH and rRNA FISH to measure associations between host and MGE and uncover the spatial context of MGE in dense biofilms.

## DISCUSSION

Here, we introduced a method for mapping MGEs in bacterial biofilms at the resolution of single cells. We optimized this method by systematically evaluating smFISH techniques to increase signal-to-noise ratio and reduce off-target binding. The resulting high sensitivity and high specificity method allowed us to map MGEs *in vitro* and in human oral plaque biofilm samples using confocal microscopy. In addition, we integrated our method with HiPR-FISH, a technique we previously created for bacterial taxon mapping in biofilms, allowing us to directly associate MGEs with their host bacteria and reveal correlations between local community structure and MGE spatial distribution. This versatile pipeline will be a valuable tool to generate and evaluate questions in microbial ecology.

Using this method, we were able to make unique observations about MGE distributions across spatial scales in *in vitro* models and human oral plaque biofilms. At the subcellular level, *in vitro*, we found that high copy plasmids without partition systems show fewer puncta than expected and localize to the poles of the cells, which supports the idea that these plasmids bunch together within the cell and do not diffuse readily in the nucleoid. We also showed that there are dramatic changes in cell shape and ribosome density associated with the number of copies of a replicating phage in *E. coli*, providing unexpected insight into the physical response of cells to infection. At the 10-100 μm scale in plaque biofilms, we demonstrated that AMR genes on plasmids and chromosomes can form clusters. We further observed clustering of two prophages at this same scale in plaque biofilms, with clusters of host cells isolated from each other by intervening non-host cells. We propose that these clustered ∼10μm regions represent spatial niches that promote short range MGE exchange in dense clusters of host taxa or support maintenance of the MGEs through replication of MGE host taxa. We also suggest that long range (∼100μm) transfer of MGEs between clusters of host taxa is limited by the need for MGEs to diffuse through the non-host biofilm. Although the literature reports that HGT is often higher in biofilms than in planktonic culture, we suggest that this observation is dependent on community spatial structure, with large variations in the local rate of HGT for a given MGE.^26,35,36^ Most importantly, we demonstrated the ability of our imaging based approach to link MGEs with their bacterial hosts including in a scenario where metagenomic sequencing could not. Our method provides the means to study the impact of taxonomic heterogeneity on the dissemination of MGEs in highly diverse natural biofilms.

We suggest that MGE mapping can serve as a direct complement for metagenomic sequencing of spatially structured microbiomes. We envision two potential application areas. First, the methods we describe could be employed to investigate the processes that govern the emergence of antibiotic resistance. Horizontal gene transfer is the predominant mechanism by which pathogens acquire antibiotic resistance, yet fundamental aspects of MGE ecology remain unknown such as the relationship between the local physical environment and the extent of MGE transfer.^26^ MGE mapping data could reveal physical parameters that influence HGT such as spatial structures or spatially clustered bacterial consortia that promote or prevent the spread of resistance elements in microbiomes. Second, MGE mapping can help address the challenge of determining bacteriophage host taxa, which is crucial given the renewed interest in phage therapy as an antibiotic alternative.^4^ In this context, MGE mapping can further be used to examine the spatial interplay between bacteria and phages in complex ecosystems, revealing the effect of local and macro structures in biofilms on phage spread, taxonomic barriers to phage infection, varying propagation modes through biofilms, the contribution of phage to biofilm structure, and biofilm “refugia” areas with reduced phage infectivity.^37,38^ These findings can then serve as a platform for developing and assessing phage therapies.

## METHODS

### Ethics statement

The protocol for volunteer recruitment and sample collection was approved by the Cornell Institutional Review Board (IRB) #2102010112.

### Human subjects sample acquisiti**on**

Volunteers were asked to refrain from cleaning their teeth for 24 hours. Volunteers then used the sharp point of a plastic toothpick to scrape the plaque from the surface of a tooth just beneath the gumline on the front and back of the tooth. They then scraped the gaps on either side of the tooth by sliding the point of the toothpick into each gap and scraping away from the gums. After each scraping action volunteers dipped the point of the toothpick into a 1.5mL sample collection tube containing 0.5ml 50% ethanol to deposit the plaque in the liquid. Samples were collected, and stored at -20°C until used.

### *E. coli* transformation and preparati**on**

Plasmid pJKR-H-TetR was acquired from addgene (https://www.addgene.org/62561/) and transformed into *E. coli* str. K-12 substr. MG1655.^8,39^ Transformed *E. coli* were streaked on LB agar Miller modification with 100 mg/L ampicillin trihydrate (MP Biomedicals, 7177-48-2) and grown overnight aerobically at 37 °C. An isolated colony was picked and grown overnight aerobically at 37 °C with 200 rpm shaking in 5 ml of LB medium Miller modification with 100 mg/L ampicillin trihydrate. 100μL overnight culture was subcultured in 10mL mod. LB with ampicillin and grown for 2hr aerobically at 37 °C with 200 rpm shaking. The culture was then split in half and one tube received 40ul 2ug/ul anhydrotetracycline (Takara, 631310) to induce GFP expression. Cultures were mixed with 10 mL 4% formaldehyde in PBS (pH 7.2 at 25 °C) and fixed for 90 minutes at room temperature. Fixed cells were pelleted (7000×g, 4 °C, 5 min.), resuspended in 500 μL cold PBS, and transferred to 1.5 mL centrifuge tubes. Cells were washed by pelleting (10000×g, 4 °C, 3 min.) and resuspended in 500 μL cold PBS, and washed again by pelleting and resuspending in 100 μL distilled water. 100 μL absolute ethanol was added to each tube to create fixed cell suspensions in 50% v/v ethanol, which were then stored at -20 °C until imaging.Wild type cells were prepared in parallel, but without ampicillin in growth media and agar.

### Phage stock preparation

*E. coli* str. K-12 substr. MG1655 was grown overnight in mod. LB medium (25 g/L Luria-Bertani broth, 300 mg/L CaCl_2_, 2 g/L D-glucose). 5 mL of overnight culture was subcultured in 50 mL mod. LB and grown aerobically at 37 °C with 200 rpm shaking for 30 minutes, then 500 μL T4 lysate was added and allowed to infect for 5 hours while shaking. Cells and cellular debris were removed from the lysate by centrifugation (7000×g, 4 °C, 10 min.) and filtration through a 0.2 μm SUPOR syringe filter (Pall). Lysate titer was determined by serially diluting lysates in mod. LB and spotting triplicate 10 μL drops of each dilution onto lawns of *E. coli* plated on mod. LB agar (mod. LB, 15 g/L agar).

### Time-course infection experiment

Replicate 7 mL mod. LB aliquots were inoculated with 100 μL overnight *E. coli* culture and grown to OD_600_ = 0.15 (∼2×10^7^ CFU/mL per growth curve analysis). High-titer T4 lysate was diluted in mod. LB and added to each culture at a multiplicity of infection of 0.01, 0.1, or 1, with uninfected cultures serving as controls. Cultures were grown aerobically at 37 °C with 200 rpm shaking. At the prescribed time points, cultures were mixed with 7 mL 4% formaldehyde in PBS (pH 7.2 at 25 °C) and fixed for 90 minutes at room temperature with continuous inversion. Fixed cells were pelleted (7000×g, 4 °C, 5 min.), resuspended in 500 μL cold PBS, and transferred to 1.5 mL centrifuge tubes. Cells were washed by pelleting (10000×g, 4 °C, 3 min.) and resuspended in 500 μL cold PBS, and washed again by pelleting and resuspending in 100 μL distilled water. 100 μL absolute ethanol was added to each tube to create fixed cell suspensions in 50% v/v ethanol, which were then stored at -20 °C until imaging. Cells were stained using Method d from *DNA-FISH protocols*.

### DNA-FISH Split-Probe design

Probes were designed using a custom Snakemake pipeline with rules written in Python using numpy and pandas.^40,41^ Target gene sequences were taken as inputs along with a reference blast database. The target was aligned to the blast database and all significant alignments were recorded for future filtering. All possible oligonucleotide probes were designed to be complementary to the coding strand of the target gene (i.e. the same sense as the mRNA) using Primer3.^42^ Pairs of Probes in this pool were identified as any probes aligning less than three base pairs distant from each other. These probe pairs were then blasted against the reference database using blastn from NCBI. On-target blast results were removed from the results using the target gene alignment IDs. Non-significant blast results were then filtered using user-defined parameters. These include maximum continuous homology (12), GC count (7), and melting temperature (46°C). All blast results with values in these parameters that were less than the specified thresholds were removed as “non-significant alignments”. The remaining blast results were considered “significant” or likely to produce off-target signal. Probe pairs were removed when both probes had off-target homologies to nearby regions in the reference database. This nearness parameter is another user-defined threshold. The remaining probe pairs were then sorted with favored probes having low levels of off-target homology. Going down the sorted list, probe pairs were then selected to tile along the gene without overlapping. Selected probes were then appended with appropriate flanking regions so that the target would be stained with the intended fluorophore (**Supp. Tab. 1**). Two base-pair spacers nucleotides between the flanking region and the probe were selected to minimize the off-target homology of the full-length probes in a similar manner to how probe pairs were sorted by blast results. The pool of selected probe pairs was then evaluated by searching for any off-target homologies where two probes were nearby each other. “Helper” probes were then selected from the Primer3 to tile along the gene without overlapping the existing probes. The final probes were then submitted for oligo synthesis to Integrated DNA Technologies (IDT) at a concentration of 200μM.

### DNA-FISH single probe design

Single probes were designed much as the split probes up to the Primer3 step. Then, instead of pairing probes, the probes were all blasted against the database and the blast results were filtered as the split probes were for “significant” off-target homologies. Probes with any significant off-target homologies were removed and the remaining probes were tiled along the target gene to ensure no overlap. The selected probes were then paired with flanking regions for the readout stain and two base pair spacers were added and optimized as in the split probe design. The resulting probes were submitted for synthesis to IDT.

### Orthogonal probe design

Probes with zero significant off-target blasts were selected from split probe pairs for different genes. For example if the left probe from a pair targeting Gene A has zero off-target blasts it is selected, then the right probe from a pair targeting Gene B is selected. The concept is that it is very unlikely these probes will hybridize close enough to each other to initiate HCR fluorescence amplification. Three right probes and three left probes were selected in this manner and pooled to create an “orthogonal” probe pool (**Supp. Tab. 1**).

### smFISH transformed *E. coli* hybridization method development protocols

Six protocols were implemented. In the first three, fixed cells suspended in 50% Ethanol were deposited on an Ultrastick slide (Electron Microscopy Sciences, 63734) and allowed to dry in a monolayer. Cells were covered in 10mg/ml Lysozyme in 10mM Tris-HCl pH 8.3, incubated at 37°C for 1hr, and washed for 2min in 1x PBS. Cells were covered with hybridization mix containing encoding probes (2x SSC, 5x denhardt’s solution, 10% Ethylene carbonate, 10% dextran sulfate, 200nM MGE probes, 200nM EUB338 probes, **Supp. Tab. 1**,**2**), incubated 4hr at 46°C, then washed for 15 min at 48°C (215mM NaCl, 20mM Tris-HCl pH7.5, 5mM EDTA). Cells were then covered with a hybridization mix containing fluorescent readout probes, incubated for 2hr at room temperature, and washed 15 min at 48°C. Slides were dried with ethanol, mountant (ThermoFisher, P36982) was deposited on the slide, a glass coverslip was placed on top, and the mountant cured for 24hr. In the first protocol only one encoding probe sequence was used with standard single fluor readout probes.^43^ In the second, ten encoding probes were used. In the third, branched readout probes were used.^11^ In the fourth protocol, hybridization chain reaction readout probes were used (prepared as previously described)^12^ at 60nM, the hybridization mix for the readout probes was altered to omit ethylene carbonate and readout was time reduced to 1.5hr. In the fifth protocol, the 10 encoding probes were substituted for 10 pairs of split encoding probes.^13^ In the fifth protocol we also added a denaturation step after removing Lysozyme from the slides. In this step we covered the cells with 50% ethylene carbonate and incubated at 60°C for 90 seconds, then immersed the slide in ice cold 70% ethanol, then ice cold 90% ethanol, then ice cold 100% ethanol for 5 minutes each. Here we also added “helper” probes to the encoding probe mix, which are unlabeled oligos with lower specificity than encoding probes that are intended to stabilize the double stranded DNA in its denatured conformation.

In the sixth protocol, we performed gelling and clearing. For this protocol, cells were deposited on 40mm round coverslips (Bioptechs, 40-1313-0319) that had been cleaned with alconox, immersed in acidic wash (5mL 37% HCl, 5mL methanol) for 30min, washed in ethanol, immersed in bind silane solution (9mL ethanol, 800μL distilled water, 100μL Bind Silane (GE, 17-1330-01), 100μL glacial acetic acid) for 30 min and allowed to air dry. Cells were then prepared as above through denaturation, then the cells were covered with Label-X solution (prepared as previously documented)^44^ and incubated for 6hr at 37°C then washed in 2x SSC for 5min, rinsed in deionized water and ethanol, and allowed to dry. The sample was covered with 50ul ice cold gel solution (4% acrylamide (1610154; Bio-Rad), 2x SSC, 0.2% ammonium persulfate (APS) (A3078; Sigma) and 0.2% N,N,N’,N’-tetramethylethylenediamine (TEMED) (T7024; Sigma) and sandwiched by a coverslip functionalized by GelSlick (Lonza; 50640).^15^ The sample was incubated at 4°C in a homemade nitrogen chamber for 1hr, then 1.5hr at 37°C. The coverslip was removed by lifting gently with tweezers from the edge, then the sample was incubated in digestion buffer (0.8 M guanidine-HCl (Sigma, G3272), 50 mM Tris·HCl pH 8, 1 mM EDTA, and 0.5% (vol/vol) Triton X-100 in nuclease-free water. 1% (vol/vol) proteinase K (New England Biolabs, P8107S)) at 100 rpm at 37°C for 12 hr, then washed in 2x SSC twice for 5min. Encoding and readout then proceeded as in the fifth protocol. Before imaging, gel samples were covered for 5min in Slowfade mountant (Thermofisher, S36963).

### Phage infection hybridization

Phage infection cells were stained using the sixth protocol from **smFISH *E. coli* method development protocols** (**Supp. Tab. 3**).

### Spectral and Airyscan Imaging

Spectral and Airyscan images were recorded on an inverted Zeiss 880 confocal microscope equipped with a 32-anode spectral detector, a Plan-Apochromat 63X/1.40 oil objective and excitation lasers at 405 nm, 488 nm, 514 nm, 561 nm, 633 nm using acquisition settings listed in **Supp. Tab. 4**. The microscope is controlled using ZEN v.2.3.

### Manual spot background filtering

Images were processed using a combination of Python scripts using numpy^40^ and interactive Jupyter notebooks to iteratively adjust and check the results of parameter adjustments. We first applied deconvolution and pixel reassignment to Airyscan images to return a super resolution image. Taking this as input, we then set a manual threshold to identify the foreground. We set the threshold such that visually distinct spots were mostly masked as separate objects. For images with high levels of non-specific signal, “blobs”, we used watershed segmentation with the background thresholded image as seed and a low intensity background thresholded image as a mask. We measured the foreground objects using skimage functions. We then removed objects larger than the threshold area. Here we set the threshold such that objects containing 1-3 neighboring spots were not removed, but objects with the continuous high signal indicative of non-specific binding were removed. We then filtered the remaining objects based on maximum intensity. Here we set the threshold to remove objects with continuous low intensity, but keep objects with high intensity peaks.

### Semi-automated image segmentation

For batches of images, an example image was selected and a zoom region within the image was selected to manually adjust segmentation parameters. In Airyscan images, segmentation parameters were set separately for cell and spot channels. In spectral images, the channels were aligned using phase cross correlation to correct for drift while switching between lasers, then the maximum projection or sum projection along the channel axis was used for segmentation. The image background mask was determined by applying a manual threshold, loading a manually adjusted background mask (as in some spot segmentation), or k-means clustering of pixel intensities. For segmentation pre-processing, images were optionally log normalized to enhance dim cells, then denoised using Chambolle total variation denoising implemented in skimage with adjustments to the weight parameter.^45,46^ In airyscan images it was sometimes necessary to blur subcellular features, so a gaussian filter could be applied with adjustments to the sigma parameter. If objects were densely packed and edge enhancement was required, we applied the local neighborhood enhancement algorithm to generate an edge-enhanced mask.^34^ In certain cases, difference of gaussians was also used for edge enhancement of the preprocessed image. We then used the watershed algorithm with peak local maxima as seeds to generate the final segmentation. Once the parameters were set, a Snakemake pipeline applied the segmentation parameters to all images in the batch. Segmented objects were measured using standard skimage functions. For spot images, local maxima were determined using skimage functions and objects with multiple local maxima were split into new objects using Pysal^47^ to generate a Voronoi diagram from the maxima to set borders between the new objects. Spots were assigned to cells based on object overlap or by radial distance between centroids.

### Spot subcellular location calculation and projection onto density map

For each spot paired with a cell, we calculated (x,y) coordinates where the x axis was the direction of the cell’s long axis and the y axis was the direction of the short axis and the magnitude of each coordinate was normalized to the average cell length and width.

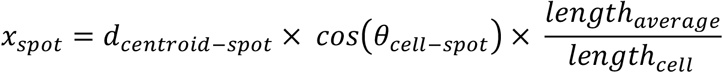

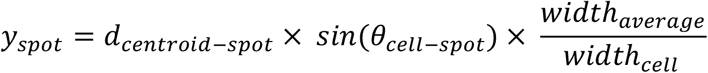

where *d*_*centroid–spot*_ is the distance between the centroid of the cell and the spot, *θ*_*cell–spot*_ is the angle between the cell’s long axis and the spot-centroid axis. We then created a grid of points to cover the average cell length and width, used the nearest neighbors algorithm to calculate the number of spots within a certain radius of each grid point, and divided by the area of the search to get a density value for each point.

### Manual Cell and spot counting

In the 30 minute and 40 minute timepoints of the phage infection, many of the infected cells had reduced 16s rRNA signal and lysed cells had caused clumps of cells to form that were difficult to segment. To count cells and classify them by their number of phage spots we used a manual counting strategy where each image was loaded into a graphic design tool (Affinity Designer) and cells of each type were counted and marked by hand. We counted a minimum of 1000 cells for each time-MOI combination.

### Prediction of phage infection rates

We used the probability mass function for a Poisson random variable to predict the fraction of cells that would encounter at least one phage

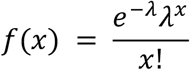

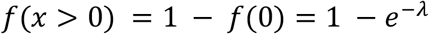

where *x* is the number of phage a cell collides with and *λ* is the ratio of average phage concentration to average cell concentration (multiplicity of infection).

### Manual seeding of transformed *E. coli* onto plaque samples

Fragments of plaque were aspirated in 50% ethanol storage solution using a 20μL pipette with a cut tip with a wide bore, deposited on a microscope slide, and allowed to dry. We then deposited 2μL of transformed *E. coli* with induced *GFP* directly on top of the plaque and allowed the slide to dry. We then proceeded through the finalized MGE-FISH method.

### Metagenomic analysis: AMR and prophage gene discovery

DNA was extracted from plaque samples using the UCP pathogen kit. The purified DNA was fragmented and prepared as an Illumina sequencing library. The samples were sequenced on an Illumina NextSeq. Raw reads were processed with PRINSEQ lite v0.20.4^48^ and trimmomatic v0.36^49^ to remove optical duplicates and sequencing adapters. Reads mapping to the human genome were discarded using BMTagger.^50^ Clean reads were assembled using SPAdes v3.14.0 (paired-end mode and –meta option)^51^ and reads were aligned to contigs using minimap2 v2.17.^52^ Contigs were resolved into metagenomic bins using vamb v3.0.2^30^ with reduced hyperparameters (-l 24, -n 384 384). Completeness and contamination of bins were evaluated with checkM v1.1.2^53^, and taxonomies were assigned to bins using GTDB-Tk v1.0.2.^31^ Read-level taxonomic relative abundance estimates were carried out with Kraken2 v2.1.2^54^ and Bracken v2.6.1.^55^ Plasmids were assembled from SPAdes assembly graphs using SCAPP v0.1^32^ using the default thresholds and scoring parameters. Lytic and lysogenic phage were identified and evaluated for induction using VIBRANT v1.2.1^28^ and PropagAtE v1.0.0,^29^ requiring a minimum length of 5000 bp and at least 10 ORFs per scaffold. Antibiotic resistance genes were annotated on contigs and mobile elements using Resistance Gene Identifier v5.2.0 against the CARD database v3.1.0 supplemented with the Resistomes & Variants dataset v3.0.8.^33^

### Plaque MGE-FISH staining

Plaque samples were stained using the fifth or sixth protocol of **smFISH *E. coli* method development protocols** with some modifications. Plaque was deposited on a microscope coverslip by aspirating 2μL of settled plaque gently from the bottom of a plaque sample collection tube with a wide bore pipette tip, depositing on the slide, and allowing excess liquid to dry. Cells were then fixed by covering with 2% formaldehyde for 10 min at room temperature, washed 5min in 1M Tris-HCl pH 7.5 for 5min, and washed in 10mM Tris-HCl pH 8.0 for 2min. Melpha X solution (prepared as previously reported)^16^ was substituted for Label X solution. Proteinase k clearing was extended to 24hr. Encoding was altered to 12hr at 46°C in a different hybridization buffer (15% formamide, 5x sodium chloride sodium citrate (SSC), 9 mM citric acid (pH 6.0), 0.1% Tween 20, 50 μg/mL heparin, 1x Denhardt’s solution, 10% dextran sulfate, 20nM encoding probes **Supp. Tab. 5-8**, 200nM EUB338 probes).^13^ After encoding, samples were washed for 5 min at 46°C in wash buffer (15% formamide, 5x SSC, 9 mM citric acid (pH 6.0), 0.1% Tween 20, 50 μg/mL heparin), 15 min at 37°C in fresh wash buffer, and 25 min at room temperature (RT) in fresh wash buffer. Readout was performed with a new readout buffer (5x SSC, 0.1% Tween 20, 10% dextran sulfate, 60nM HCR hairpins, 200nM EUB338 readout probes). After readout, samples were washed for 5min at RT in 5x SSCT (5x SSC, 0.1% Tween 20), 30 min at RT in fresh 5x SSCT twice more, then 5min in fresh 5x SSCT. Samples were covered with Slowfade mountant before imaging.

### Spatial autocorrelation analysis

A neighbor spatial connectivity matrix was constructed from cell segmentation centroids using a Voronoi diagram algorithm from Pysal Each cell was given a binary mark indicating presence of MGE spot. The weight matrix and marked cells were used in a global Moran’s I test from Pysal to calculate spot autocorrelation. The measured Moran’s I value was compared against a simulation based null model that spots are randomly distributed within the cell space.

### Large scale spot density analysis

After spot segmentation, the universal 16s rRNA signal was used to create a global mask to identify the foreground. For each pixel in the foreground, we used the nearest neighbors algorithm to calculate the number of spots within a certain radius of each grid point, and divided by the area of the search to get a density value for each point.

### Spatial association measurements

We performed two versions of spot colocalization. First in a given color channel, for each spot we used the nearest neighbors algorithm to determine whether there were spots of the other color(s) within a 0.5μm radius and calculated the fraction of spots colocalized with each of the other colors based on the number of spots in the reference channel. We repeated the measurement for each color channel. In the second version, we overlaid the spots from each channel (labeled as different spot types), divided the image into a grid of squares with 5μm edges, classified each square based on the number of spot types present, counted the number of squares of each type, and normalized by the total number of squares with at least one spot type.

### Genus level probe design

We performed full length 16s rRNA sequencing and taxonomic classification as previously described^34^ on the extracted DNA used for metagenomic sequencing in **Metagenomic analysis: AMR and prophage gene discovery**. We searched for previously designed genus level FISH probe sequences^27^ and blasted the probes against our full length 16s rRNA data using blastn. We filtered results to remove “non-significant” alignments as defined above in **DNA-FISH Split-Probe design**, determined the fraction of significant alignments to non-target genera, and removed probes with off-target rate greater than 0.1. We then selected 5-bit binary barcodes for each genus such that most barcodes were separated by a hamming distance of 2. Based on the binary barcodes we concatenated a readout sequence to the three prime end of each probe sequence such that the readout sequence would hybridize the appropriate fluorescent readout probe for the barcode (**Supp. Tab. 9**). For barcodes with multiple colors in the barcode, we created separate probes concatenated with each readout sequence. We created barcodes that used only the 488 nm, 514 nm, and 561 nm lasers, thus reserving the 633 nm laser for MGE-FISH and the 405 nm laser for the universal EUB338 16s rRNA stain. For stains where we targeted only 5 genera, we simply used a different fluorophore for each genus probe.

### Combined MGE-FISH and HiPR-FISH staining

Samples were prepared with the sixth protocol in “smFISH transformed *E. coli* hybridization method development protocols” and as in “Plaque MGE-FISH staining” except for the hybridization buffer, which included 20nM of pooled genus probes, and the readout buffer which included 200nM of each of the five fluorescent readout probes.

### Pixel level spectral deconvolution and taxon assignment

We aligned the laser channels of the spectral images using phase cross correlation, then we performed gaussian blurring (sigma=3) on each spectral channel to reduce the noise in each pixel’s spectra. We acquired a maximum intensity projection along the channel axis, selected a background threshold, and generated a mask. To account for nonspecific binding, which generates a low intensity background signal with the “11111” (all 5 fluorophores) spectral barcode, we multiplied the “11111” reference spectrum by a scalar and subtracted the scaled spectrum from each pixel’s measured spectrum (reference spectra for each barcode were collected as previously described)^34^. We visualized the pixel spectra before and after subtraction and adjusted the scalar such that the visually apparent background was removed (scalar=0.05). The adjusted pixel spectra were stored in a “pixel spectra matrix” with the following shape: (*number of pixels, number of spectral channels*). The reference spectra for all barcodes were sum normalized and merged in a “reference spectra matrix” with the following shape: (*number of spectral channels, number of barcodes*). We performed matrix multiplication between the “pixel spectra matrix” and the “reference spectra matrix” to get a “classification matrix” with shape: (*number of pixels, number of barcodes*). Separately, we evaluated the reference spectra and created a boolean array indicating whether or not we expected a signal from each of the three lasers. We merged these arrays into a “reference laser presence” matrix with shape: (*number of lasers, number of barcodes*). Then, for each adjusted pixel spectrum we measured the maximum value for each laser, normalized these values by the highest of the three values, and set minimum threshold values (threshold_488_=0.3, threshold_514_=0.4, threshold_561_=0.3) to create a “pixel laser presence” boolean matrix with shape: (*number of pixels, number of lasers*). We performed matrix multiplication between the “pixel laser presence” matrix and the “reference laser presence” matrix to get a matrix with shape: (*number of pixels, number of barcodes*). We performed element-wise multiplication between this matrix and the “classification matrix” to remove barcodes from the classification matrix if the signal from one of the lasers was too low. For each pixel, we selected the barcode with the highest value in the adjusted “classification matrix”.

### Cell segmentation and taxon assignment

For each object in the cell segmentation, if all the pixels within the object were assigned to the same taxon, we assigned that taxon to the object. If multiple taxa were represented in the cell pixels, the object was split into multiple new objects such that each new object encompassed pixels of only one taxon.

### Taxon-spot spatial association measurements

We created a subset of the cell centroids for each taxon. Then for each taxon we used the nearest neighbor algorithm to measure the distance from each spot to the nearest cell of that taxon and counted the number of spots where distance was less than 0.5μm. To calculate the fraction of spots and taxon cells, we divided the count by the total number of spots and total number of taxon cells respectively.

### Random simulation of spot distribution

We used the foreground mask to create a list of pixel coordinates within the plaque cells, then used a random integer generator to select pixels by their list index. We used the randomly selected pixel coordinates as simulated spots and counted taxon-spot spatial associations as described above. This was repeated for 1000 simulations and we calculated the mean and standard deviation for the count values for each taxon. We then calculated the z-score for the count values: *z =* (*count − mean*) / *standard deviation*.

## Author Contributions

B.G., H.S., and I.D.V. conceived the study. B.G., H.S., O.F., Y.N., A.V., P.D., W.R.Z., I.B., and I.D.V. designed staining and imaging methods and validation experiments. B.G., O.F., and Y.N. performed staining and imaging methods and validation experiments. B.G. and H.S. collected volunteer samples. A.V. analyzed metagenomic sequencing data to identify target genes and designed and performed the *in vitro* phage infection system. P.D. designed and performed the *in vitro* plasmid system. B.G., H.S., and O.F. wrote the probe design and image analysis pipelines. B.M.G. and I.D.V. wrote the manuscript and prepared the figures. All authors read and edited the manuscript.

## Competing interests

H.S. is a co-founder at Kanvas Biosciences. I.D.V. is a member of the Scientific Advisory Board of Karius Inc., and GenDX and co-founder of Kanvas Biosciences. H.S. and I.D.V. are listed as inventors on patents related to multiplexed imaging methods.

## Acknowledgements

We thank R. M. Williams and J. M. Dela Cruz for assistance with microscopy; T. Doerr for providing materials; and M. Mantri, D. W. McKellar, J. Jones, L. Takayasu, S. Arias, and T. Ciavatti for discussions and feedback. This work was supported by an instrumentation grant from the Kavli Institute at Cornell and by US National Institutes of Health (NIH) grants 1DP2AI138242 to I.D.V. and 1R33CA235302 to I.D.V., W.Z. and I.L.B. Imaging data were acquired in the Cornell Biotechnology Resource Center Imaging Facility using the shared, NYSTEM (CO29155)- and NIH (S10OD018516)-funded Zeiss LSM880 confocal and multiphoton microscope.

## Code Availability

The specific implementation of code to generate figures presented here is available on GitHub at https://github.com/benjamingrodner/hipr_mge_fish. The generalized pipeline for segmentation is available at https://github.com/benjamingrodner/pipeline_segmentation, while the generalized implementation of probe design is available at https://github.com/benjamingrodner/FISH_split_probe_design.

## Data Availability

Illumina and PacBIO sequencing data are available at the NCBI Sequence Read Archive (SRA) with accession number <u>PRJNA981198</u>. Microscopy data have been deposited to Zenodo at https://doi.org/10.5281/zenodo.8015720 (**Fig. 1b, Fig. S1a**,**b**), https://doi.org/10.5281/zenodo.8015754 (**Fig. 1c, Fig. S1c**,**d**), https://doi.org/10.5281/zenodo.8015832 (**Fig. 2, Fig. S2, 3**), https://doi.org/10.5281/zenodo.8016062 (**Fig. 3, 4**).

