## Supplementary Figures for "Spatial Mapping of Mobile Genetic Elements and their Cognate Hosts in Complex Microbiomes"

**Code repository:** https://github.com/benjamingrodner/hipr_mge_fish

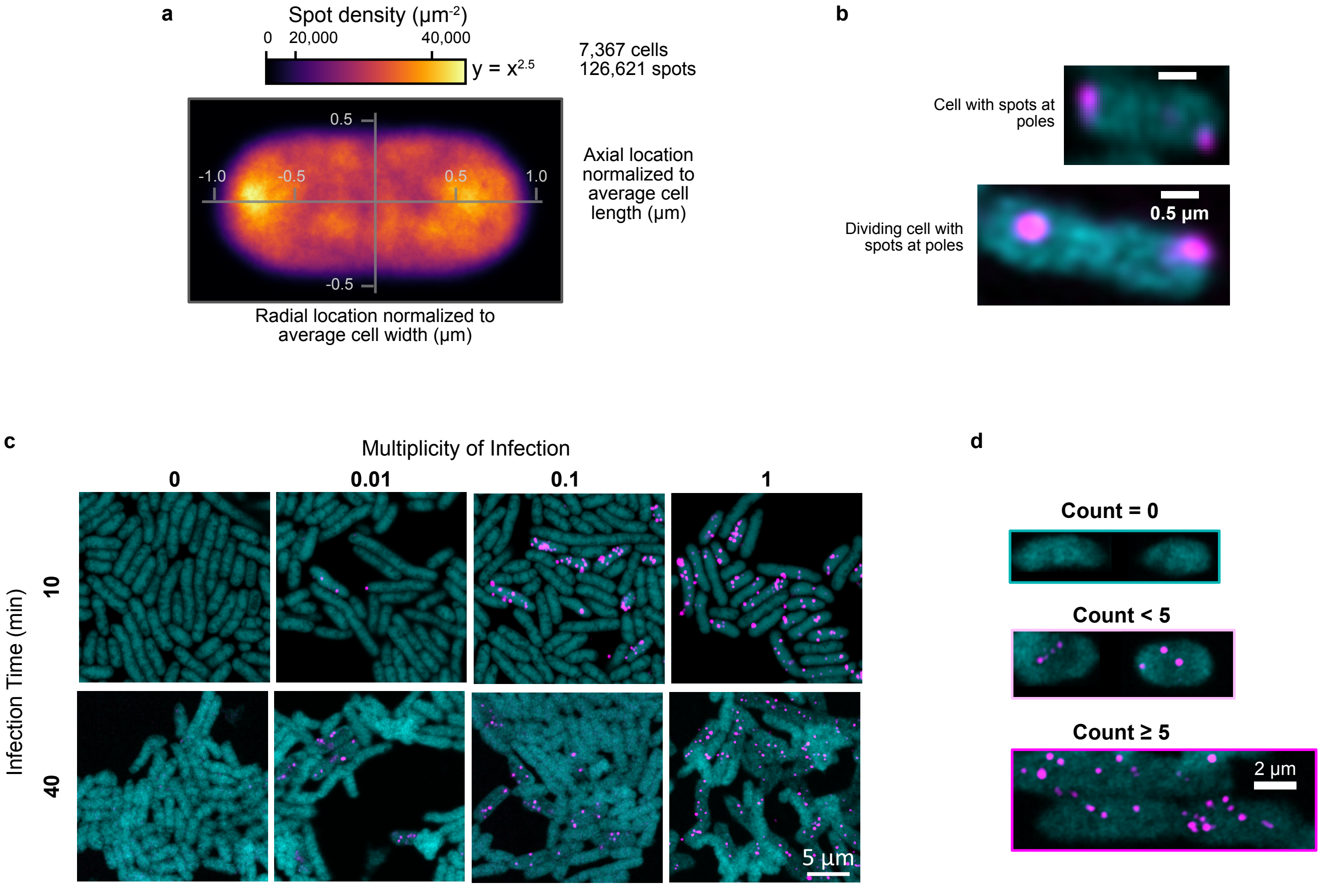


**Figure S1. In vitro single MGE-FISH on plasmids and phage. a** Subcellular spot locations MGE-FISH staining of *GFP*-plasmid normalized to average cell shape plotted as spot density. The colormap is nonlinear and maps to values using a power law where color value $x$ maps to density value $y=x^{2.5}$. **b** Example zoomed raw images of cells where spots are located at the poles. **c** Example images of MGE-FISH staining of T4 phage *gp34* gene in T4 phage infection time series. Cyan: 16s rRNA, magenta: GFP DNA. **d** Example cells showing examples for the different classification of cells in the manual counting data in **Fig. 1c**.

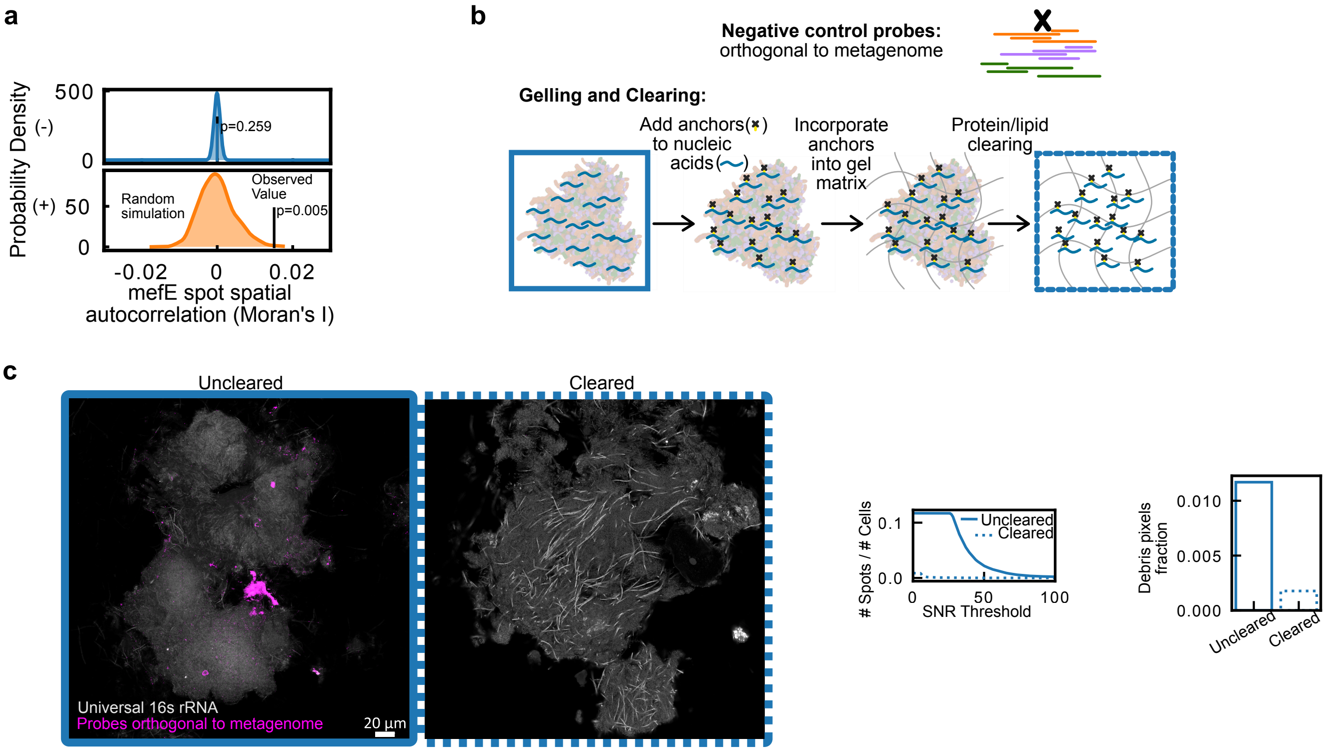


**Figure S2. Evaluation of MGE-FISH in plaque biofilms. a** Spatial autocorrelation of mefE spots using Moran's I statistic. Colored vertical bar indicates the observed value, black vertical bar indicates the mean value of the simulation, and the shaded area indicates the histogram of the simulation. For the simulation spots were randomly redistributed on the same set of cell segmentations 1000 times. **b** *Top*: diagram of orthogonal control probes that should produce no signal. *Bottom*: diagram of the gel embedding, nucleic acid anchoring, and sample clearing process. **c** *Left*: Example images showing the off target signal from orthogonal probes in uncleared and cleared plaque samples. *Center*: spot counts normalized by number of cells as a function of signal to noise ratio (SNR). *Right*: Measurement of non-spot pixels normalized by cell pixels.

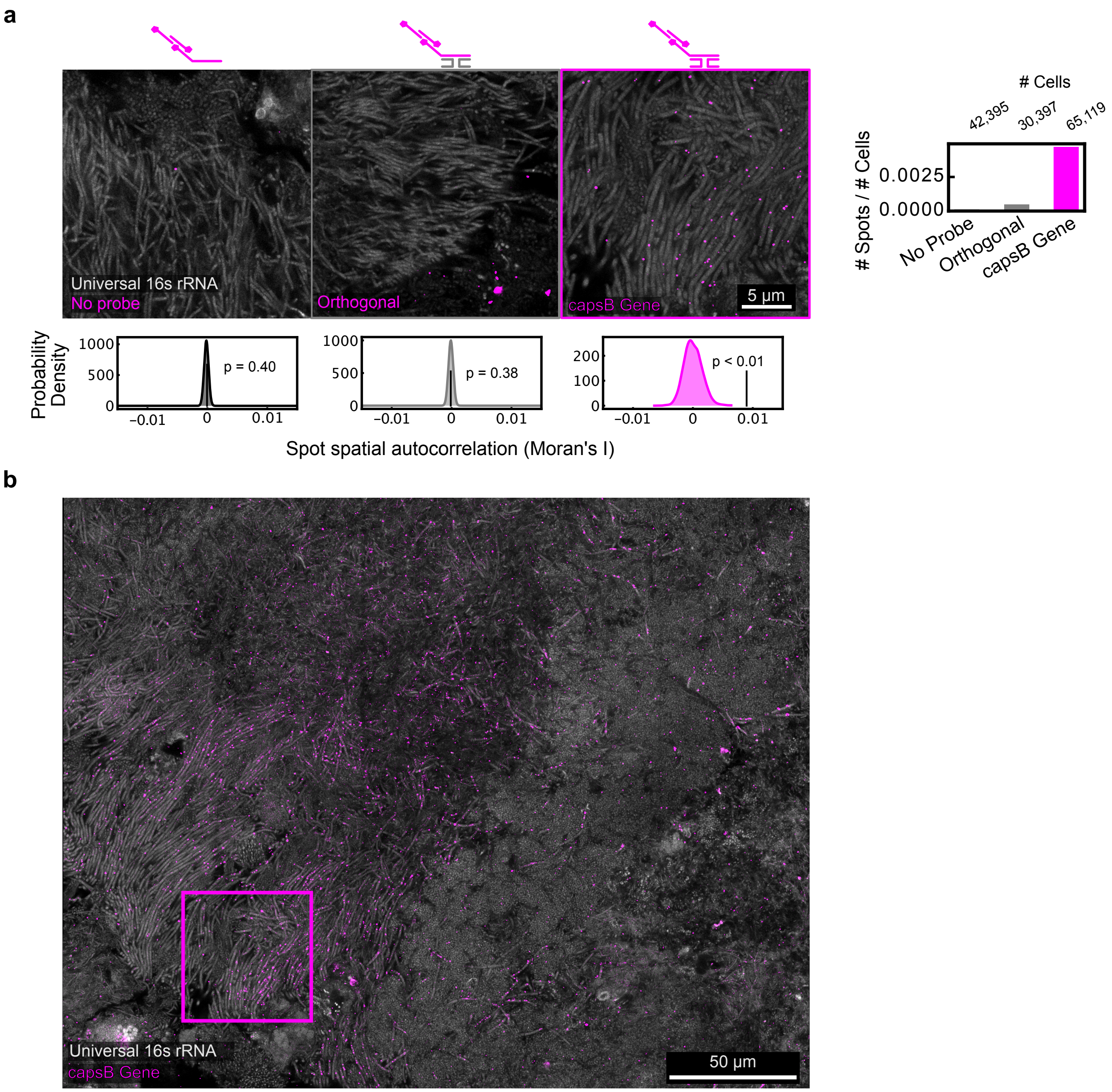


**Figure S3. Control experiments for T7-like prophage capsB minor capsid protein in Plaque. a** *Top left*: example images showing regions of cells with similar morphology cells. From right to left the images are: MGE-FISH controls with no encoding probes, encoding probes that are orthogonal as determined by metagenomic analysis, or probes targeting *capsB*. *Top right*: spot counts from each control normalized by number of cells. *Bottom*: observed Moran's I spatial autocorrelation values (vertical black lines) compared to 999 simulations of random spot distribution (filled curves). **b** Example FOV showing a large hotspot of prophage (~100µm). Inset square shows the location of the *capsB* example image in **a**.
